# Peroxiredoxin-mediated HMGB1 oxidation and secretion in response to inflammatory stimuli

**DOI:** 10.1101/258285

**Authors:** Man Sup Kwak, Hee Sue Kim, Khulan Lkhamsuren, Young Hun Kim, Myung Gil Hahn, Jae Min Shin, In Ho Park, Se Kyung Lee, Sue Goo Rhee, Jeon-Soo Shin

**Affiliations:** Department of Microbiology, Yonsei University College of Medicine, Seoul 03722, Korea; Brain Korea 21 PLUS Project for Medical Science, Yonsei University College of Medicine, Seoul 03722, Korea; Severance Biomedical Science Institute, Yonsei University College of Medicine, Seoul 03722, Korea; Institute for Immunology and Immunological Diseases, Yonsei University College of Medicine, Seoul 03722, Korea; Center for Nanomedicine, Institute for Basic Science (IBS), Yonsei University, Seoul 03722, Korea

## Abstract

The nuclear protein HMGB1 (high mobility group box 1) is secreted by monocytes-macrophages in response to inflammatory stimuli and serves as a danger-associated molecular pattern. Acetylation and phosphorylation of HMGB1 are implicated in the regulation of its nucleocytoplasmic translocation for secretion, although inflammatory stimuli are also known to induce H_2_O_2_ production. Here we show that H_2_O_2_-induced oxidation of HMGB1 that results in formation of an intramolecular disulphide bond between Cys^23^ and Cys^45^ is necessary and sufficient for its nucleocytoplasmic translocation and secretion. The oxidation is catalysed by peroxiredoxin I (PrxI) and PrxII, which are first oxidized by H_2_O_2_ and then transfer their disulphide oxidation state to HMGB1. The disulphide form of HMGB1 showed a higher affinity for the nuclear exportin CRM1 compared with the reduced form. Lipopolysaccharide (LPS)–induced HMGB1 secretion was greatly attenuated in macrophages derived from PrxI or PrxII knockout mice, as was the LPS-induced increase in serum HMGB1 levels in these mice.

## Introduction

High mobility group box 1 (HMGB1) is a nonhistone nuclear protein that functions as a DNA chaperone during chromatin reorganization and transcriptional regulation (Lee et al, 2014; Lotze & Tracey, 2005) and which shuttles between the nucleus and cytoplasm of mammalian cells. Exit of HMGB1 from the nucleus occurs by both passive diffusion and an actively controlled pathway mediated by chromosome region maintenance protein 1 (CRM1), with import of the protein back into the nucleus also being actively controlled (Kang et al, 2014). The trafficking of HMGB1 between the nucleus and cytoplasm has been thought to be regulated by protein acetylation and phosphorylation (Bonaldi et al, 2003; Youn & Shin, 2006). When HMGB1 is hypoacetylated or hypophosphorylated, the rate of nuclear import exceeds that of export, resulting in a predominantly nuclear localization. When HMGB1 is acetylated or phosphorylated at its nuclear localization signal (NLS) sequences in the nucleus, however, it is still able to exit the nucleus but reimport is inhibited, resulting in its accumulation in the cytoplasm. HMGB1 is passively released from cells via pyroptosis and necroptosis, but it is also actively secreted from monocytes and macrophages in response to exposure of the cells to inflammatory stimuli such as lipopolysaccharide (LPS) and tumor necrosis factor α (TNFα). HMGB1 does not contain a leader sequence, with such secretion occurring through a vesicle-mediated pathway (Gardella et al, 2002).

Secreted HMGB1 acts as a damage-associated molecular pattern (DAMP) molecule to signal danger to surrounding cells, triggers inflammation, and activates innate and adaptive immunity by binding to a plethora of cell surface receptors including Toll-like receptor 2 (TLR2), TLR4, TLR9, receptor for advanced glycation end products (RAGE), and C-X-C chemokine receptor 4 (CXCR4). HMGB1 also interacts with the pathogen-associated molecular pattern (PAMP) molecules LPS and lipoteichoic acid (Kwak et al, 2015; Youn et al, 2008) as well as with endogenous C-X-C motif chemokine 12 (CXCL12) (Schiraldi et al, 2012), interleukin-1β (Sha et al, 2008), and nucleosomes (Tian et al, 2007) to augment inflammation. Furthermore, HMGB1 mediates autophagosome formation by binding to beclin-1 (Tang et al, 2010), and it inhibits polyglutamine aggregate formation in the cytoplasm through a chaperone-like function (Min et al, 2013).

HMGB1 contains three conserved cysteine residues: Cys^23^, Cys^45^, and Cys^106^. The redox state of these cysteines gives rise to at least three forms of HMGB1 that are designated “reduced HMGB1” (all three cysteine residues in the thiol state), “disulphide HMGB1” (an intramolecular disulphide bond between Cys^23^ and Cys^45^; Cys^106^ in the thiol state), and “sulfonic HMGB1” (all three cysteines in the hyperoxidized sulfonic acid state). The specific functions of HMGB1 have recently been shown to be dependent on the redox state of its cysteines (Venereau et al, 2012), with reduced HMGB1 forming a complex with CXCL12 to promote the migration of immune cells (Venereau et al, 2012), disulphide HMGB1 (but not reduced HMGB1) interacting with a TLR4 adaptor protein to activate TLR4 and induce proinflammatory responses (Yang et al, 2010), and sulfonic HMGB1 manifesting neither cytokine-nor chemokine-like function (Kazama et al, 2008). In addition, displacement of histone H1 from DNA is promoted by reduced HMGB1, whereas the binding of histone H1 to chromatin is dependent on HMGB1 oxidation (Kohlstaedt et al, 1987; Polanska et al, 2014). The redox state of HMGB1 in the nucleus thus plays an important role in transcriptional regulation.

HMGB1 present inside cells (nucleus and cytoplasm) is predominantly in the fully reduced state, whereas the presence of disulphide HMGB1 in serum has been associated with inflammation-related pathologies(Antoine et al, 2012; Liesz et al, 2015; Lundback et al, 2016; Palmblad et al, 2015). However, fundamental issues regarding the oxidation of Cys^23^ and Cys^45^ of HMGB1—including the mechanism by which such oxidation occurs in response to inflammatory signals, the subcellular compartment where the oxidation occurs, and the effect of oxidation on HMGB1 secretion—have remained unclear.

Hydrogen peroxide (H_2_O_2_) is produced by cells in response to various extracellular stimuli and propagates intracellular signaling by oxidizing protein thiols (Rhee et al, 2017). Cells are also equipped with H_2_O_2_-eliminating enzymes such as catalase, glutathione peroxidase, and peroxiredoxin (Prx). The Prx family of peroxidases reduces H_2_O_2_ or alkyl peroxide, with the thiol of a conserved cysteine residue (the peroxidatic Cys, or C_P_–SH) serving as the site of oxidation by peroxides. In addition to C_P_–SH located in the NH_2-_terminal region of the molecule, most Prx enzymes contain an additional conserved cysteine (the resolving Cys, or C_R_–SH) in the COOH-terminal region. Mammalian cells express six isoforms of Prx (PrxI to PrxVI) that differ in their subcellular localization (Wood et al, 2003). PrxI and PrxII are thus localized in the cytosol and nucleus, whereas PrxIII and PrxIV are restricted to mitochondria and the endoplasmic reticulum, respectively. During the catalytic cycle of Prx enzymes, C_P_–SH is first oxidized by H_2_O_2_ to sulfenic acid (C_P_–SOH), which then reacts with C_R_–SH to form a disulphide that is subsequently reduced by an appropriate electron donor to complete the cycle.

Peroxiredoxins are abundant proteins, with PrxI and PrxII together constituting a total of 0.2% to 1% of soluble protein in cultured mammalian cells (Chae et al, 1999). In addition, Prxs possess an active-site pocket that gives rise to a high-affinity (submicromolar dissociation constant) peroxide binding site that is lacking in catalase and glutathione peroxidase (Rhee & Kil, 2017). This pocket is arranged so as to elicit rapid oxidation of C_P_– SH by H_2_O_2_, with the second-order rate constant for this oxidation being several orders of magnitude greater than that for the oxidation of thiols in well-characterized H_2_O_2_ target proteins. It was thus at first unclear how H_2_O_2_ is able to oxidize specific cysteine thiols of its target proteins in the presence of such abundant and efficient antioxidant enzymes. Two mechanisms to overcome this kinetic disadvantage with regard to oxidation by H_2_O_2_ have been proposed. One mechanism is transient localized inactivation of Prxs so as to protect H_2_O_2_ from destruction, which allows the accumulation of H_2_O_2_ in specific regions of the cell without global redox disturbance. The second mechanism is oxidation of PrxI or PrxII by H_2_O_2_ and subsequent transfer of the oxidation state to the target protein. Such an H_2_O_2_ sensor and transducer function of Prxs has been demonstrated in the oxidation of several proteins including protein disulphide isomerase (PDI) (Kakihana et al, 2013), apoptosis signal– regulating kinase 1 (Zhang et al, 2015), mitogen-activated protein kinase (MAPK) (Latimer & Veal, 2016; Turner-Ivey et al, 2013), and the transcription factor STAT3 (Sobotta et al,2015).

We now show that disulphide formation between Cys^23^ and Cys^45^ of HMGB1 is mediated by PrxI and PrxII in the nucleus and is essential for the nucleocytoplasmic translocation and secretion of HMGB1. Inflammatory stimuli such as LPS and TNFα as well as chemicals including phorbol 12-myristate 13-acetate (PMA) and trichostatin A (TSA) have been shown to induce the phosphorylation or acetylation of HMGB1 at its NLS sequences and thereby to promote its nucleocytoplasmic translocation for secretion (Bonaldi et al, 2003; Youn & Shin, 2006). These inflammatory stimuli and chemicals also induce the intracellular production of H_2_O_2_. We here show that H_2_O_2_ is first sensed by PrxI and PrxII, which then transfer the oxidant signal to HMGB1, resulting in formation of the Cys^23^–Cys^45^ disulphide bond. Importantly, whereas such disulphide formation is sufficient for HMGB1 secretion,posttranslational modification of the NLS sequences is not essential for secretion but rather appears to function cooperatively with disulphide formation in this regard. Finally, we found that the LPS-induced increase in the circulating concentration of HMGB1 was significantly attenuated in mice lacking the PrxI or PrxII gene.

## Results

### HMGB1 oxidation mediated by PrxI and PrxII in the nucleus

LPS has been shown to induce HMGB1 secretion by promoting the acetylation of HMGB1 in the nucleus. Ligation of TLR4 by LPS also elicits H_2_O_2_ production through activation of NADPH oxidase 4 (Nox4) (Park et al, 2004). To examine the effect of LPS on the formation of disulphide HMGB1, we stimulated mouse bone marrow–derived macrophages (BMDMs) with LPS and then subjected the nuclear fraction of these cells to nonreducing SDS– polyacrylamide gel electrophoresis (PAGE) followed by immunoblot analysis. Disulphide HMGB1 is more compact and thereby migrates faster in the electrophoretic gel compared with reduced HMGB1. HMGB1 in the nucleus of unstimulated cells existed predominantly in the fully reduced form (Fig 1A). Stimulation with LPS, however, induced a time-dependent increase in the nuclear abundance of disulphide HMGB1 (Fig 1A).

**Figure 1.**
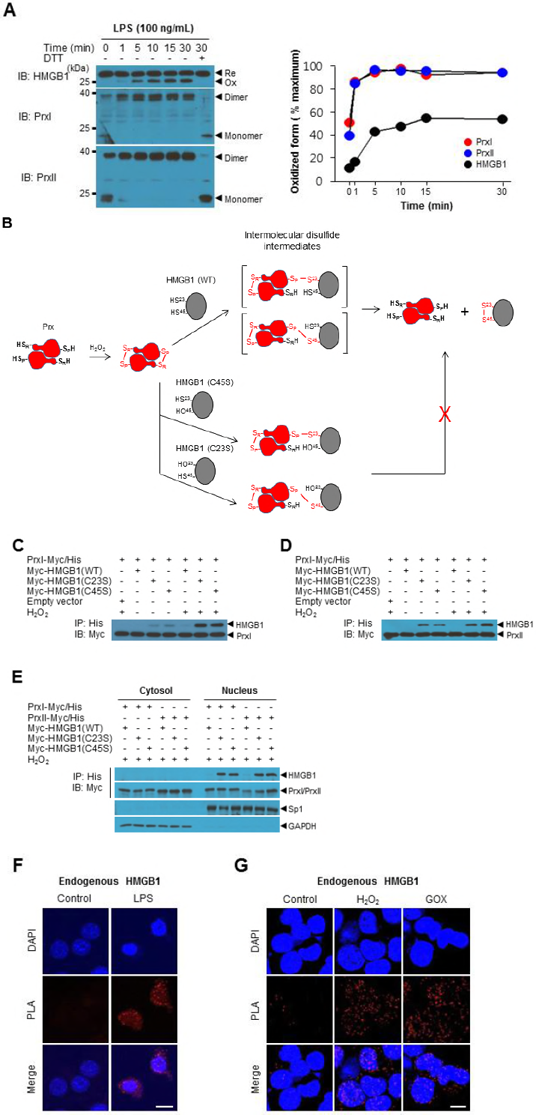
HMGB1 oxidation mediated by PrxI and PrxII in the nucleus. A. Mouse BMDMs were treated with LPS (100 ng/ml) for various times, after which the nuclear fraction of cell lysates was isolated and subjected to nonreducing SDS-PAGE followed by immunoblot (IB) analysis with antibodies to HMGB1, to PrxI, or to PrxII (left panel). The nuclear fraction of cells stimulated with LPS for 30 min was also exposed to 5 mM DTT before analysis as a control. The intensities of bands corresponding to reduced (Re) and oxidized (Ox) forms of HMGB1 and to reduced (monomeric) and oxidized (dimeric) forms of PrxI and PrxII were determined as mean values from two independent experiments (right panel). B. Mechanistic scheme for the formation of disulphide HMGB1 catalysed by PrxI or PrxII. PrxI or PrxII is first oxidized by H_2_O_2_, resulting in the formation of a homodimer linked by intermolecular disulphides between C_P_ and C_R_. The disulphide oxidation state of Prx is then transferred to reduced HMGB1, resulting in the formation of disulphide HMGB1 with an intramolecular linkage between Cys^23^ and Cys^45^. Resolution of the intermolecular disulphide intermediate is not possible in the case of C23S or C45S mutants of HMGB1. C, D HEK293T cells that had been transfected with expression vectors for Myc epitope– tagged human HMGB1(WT), HMGB1(C23S) or HMGB1(C45S) or with the corresponding empty vector as well as with those for Myc- and His_6_-tagged human PrxI (C) or PrxII (D) were incubated in the absence or presence of 50 μM H_2_O_2_ for 30 min, after which whole cell lysates were subjected to immunoprecipitation (IP) with antibodies to the His_6_ tag and the resulting precipitates were subjected to immunoblot analysis with antibodies to the Myc tag. E. The whole cell lysates prepared in (C) and (D) were also separated into cytosolic and nuclear fractions before immunoprecipitation and immunoblot analysis. The fractions were also directly subjected to immunoblot analysis of Sp1 and GAPDH as marker proteins for the nucleus and cytosol, respectively. F, G PLA assay for detection of the interaction between endogenous HMGB1 and PrxI. MEFs were incubated in the absence (Control) or presence of LPS (1 μg/ml) for 1 h (F) or 50 μM H_2_O_2_ or GOX (5 mU/ml) for 30 min (G). The cells were then fixed and incubated with antibodies to HMGB1 and to PrxII before the addition of PLA probes and processing for detection of the intermolecular interaction (red fluorescence) by confocal microscopy. Nuclei were stained with 4',6-diamidino-2-phenylindole (DAPI, blue fluorescence). Scale bars, 10 üm. See also Expanded View figure 1.

We examined the possibility that nuclear PrxI or PrxII might function as an H_2_O_2_ sensor and transducer for the formation of disulphide HMGB1. PrxI and PrxII are obligate homodimers arranged in a head-to-tail manner. In the presence of H_2_O_2_, the C_P_–SH residue (Cys^52^ for PrxI and Cys^51^ for PrxII) is oxidized to sulfenic acid (C_P_–SOH), which then reacts with the C_R_–SH residue (Cys^173^ for PrxI and Cys^172^ for PrxII) of the other subunit to form a disulphide-linked dimer (Fig 1B). Nonreducing SDS-PAGE and immunoblot analysis revealed that LPS stimulation rapidly increased the abundance of the disulphide-linked dimeric forms of both PrxI and PrxII and that this effect preceded the oxidation of HMGB1 (Fig 1A), suggesting that PrxI/II might indeed mediate the oxidation of HMGB1 in the nucleus. As expected, neither the higher-mobility form of HMGB1 nor dimeric PrxI/II was observed if dithiothreitol (DTT) was added to the nuclear extracts before electrophoresis (Fig 1A).

The putative Prx-mediated oxidation of HMGB1 would likely be initiated by attack of the disulphide of the Prx dimer by either Cys^23^–SH or Cys^45^–SH of HMGB1, resulting in the formation of a transient intermolecular disulphide–linked intermediate containing both Prx and HMGB1 (Fig 1B). Resolution of the intermolecular disulphide by either Cys^23^–SH or

Cys^45^–SH of HMGB1, whichever does not participate in the intermolecular disulphide linkage, would then result in regeneration of reduced Prx and formation of disulphide HMGB1 (Fig 1B). To verify this possibility by detection of the disulphide-linked HMGB1-Prx complex, we transfected human embryonic kidney (HEK) 293T cells with expression vectors for Myc epitope–tagged wild-type (WT) or Cys^23^-to-Ser (C23S) or Cys^45^-to-Ser (C45S) mutant forms of HMGB1 as well as for Myc- and His_6_-tagged forms of PrxI or PrxII.The cells were then stimulated with H_2_O_2_, and the amount of Myc-HMGB1 that coimmunoprecipitated with Myc/His_6_-tagged PrxI or PrxII from cell lysates was determined by immunoblot analysis. Coprecipitation of Myc-HMGB1(C23S) and Myc-HMGB1(C45S) with Myc/His_6_-tagged PrxI or PrxII was apparent in the absence of H_2_O_2_, and stimulation of the cells with H_2_O_2_ greatly increased the extent of such coprecipitation (Fig 1C and D).Preparation of nuclear and cytosolic fractions from the cell lysates revealed that the coprecipitation was apparent with the former fraction but not the latter (Fig 1E). These results thus indicated that PrxI/II-mediated generation of disulphide HMGB1 occurs via formation of an intermolecular disulphide in the nucleus as proposed in Figure 1B. The disulphide-linked complexes of HMGB1 with either PrxI or PrxII were also directly detected by analysis of the cell lysates by nonreducing SDS-PAGE (Fig EV1A). The essential roles of the two cysteine residues of HMGB1 (Cys^23^ and Cys^45^) and the two cysteines of PrxI/II (C_P_ and C_R_) information of the intermolecular disulphide–linked intermediates were demonstrated by analysis of HEK293T cells expressing a double Cys-to-Ser mutant of either HMGB1[HMGB1(C23/45S)] or Prx [PrxI(C52/173S) or PrxII(C51/172S)] (Fig EV1B). We also detected DTT-sensitive co-immunoprecipitation of purified PrxI or PrxII with purified HMGB1 in the presence of H_2_O_2_ in vitro (Fig EV1C). The amounts of PrxI and PrxII in the nuclear fraction of HEK293T cells, Raw264.7 mouse macrophages, and mouse embryonic fibroblasts (MEFs) were similar and in the range of 60 to 90 pg of each protein per microgram of total nuclear protein (Fig EV1D).

We performed a proximity ligation assay (PLA) in MEFs to examine formation of the disulphide between endogenous HMGB1 and PrxII proteins. LPS stimulation induced a marked increase in the number of PLA signals in the nucleus of MEFs, indicative of a substantial extent of complex formation (Fig 1F). The number of PLA signals due to the HMGB1-PrxII intermediate was also greatly increased by exposure of the cells to H_2_O_2_ or glucose oxidase (GOX) (Fig 1G). We also performed a PLA assay with HEK293T cells expressing Myc epitope–tagged WT, C23S, or C45S forms of HMGB1 in order to detect formation of disulphide-linked complexes of these proteins with endogenous PrxII. Treatment of the cells with H_2_O_2_ or GOX increased the number of PLA signals in the nucleus of cells expressing each form of HMGB1 (Fig EV1E-G). As expected, however, the number of PLA signals was greater in the cells expressing HMGB1(C23S) or HMGB1(C45S) than in those expressing HMGB1(WT) (Fig EV1E-G).

We next evaluated the effect of the absence of PrxII on the formation of disulphide HMGB1 with the use of immortalized WT and PrxII-deficient (KO) MEFs. The protein kinase C (PKC) activator PMA is thought to stimulate HMGB1 secretion by inducing HMGB1 phosphorylation at serine residues located in the NLS sequences (Oh et al, 2009; Youn & Shin, 2006). Like LPS, PMA also induces H_2_O_2_ production by activating Nox enzymes (Karlsson et al, 2000). Stimulation with LPS or PMA increased the abundance of disulphide HMGB1 in WT and PrxII KO MEFs, but this effect was markedly attenuated in the latter cells compared with the former for both stimulants (Fig 2A and B). The amount of disulphide HMGB1 produced in PrxII KO MEFs in response to GOX or H_2_O_2_ treatment was increased greatly by ectopic expression of PrxII(WT) but was not affected by that of PrxII(C172S) (Fig 2C and D). The C_P_ residue (Cys^51^) of PrxII(C172S) is still sensitive to H_2_O_2_ and is readily oxidized to sulfenic acid (C_P_–SOH). The failure of PrxII(C172S) to mediate HMGB1 oxidation in PrxII KO MEFs suggests that PrxII transfers its oxidation state to HGMB1 via the disulphide-linked dimer but not via the sulfenic form (C_P_–SOH). Similar to the results obtained with PrxII KO MEFs, we found that HMGB1 oxidation induced by GOX or H_2_O_2_ in PrxI KO MEFs was greatly enhanced by ectopic expression of PrxI(WT) but not by that of PrxI(C173S) (Fig EV2).

**Figure 2.**
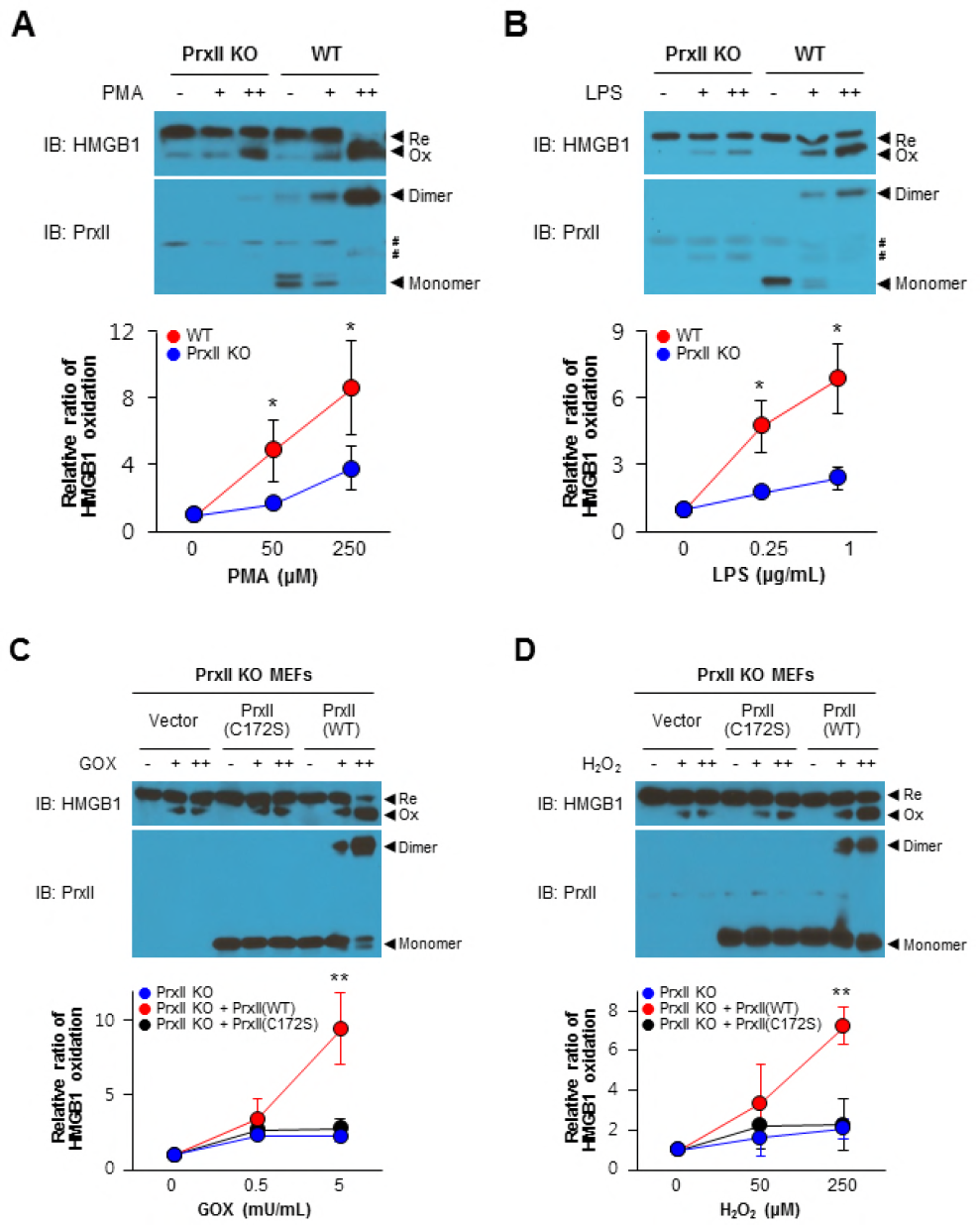
Ablation of PrxII impedes formation of disulphide HMGB1. A, B Immortalized WT or PrxII KO MEFs were incubated for 30 min in the absence or presence of PMA (50 or 250 nM) (A) or LPS (0.25 or 1 μg/ml) (B), after which cell lysates were subjected to nonreducing SDS-PAGE followed by immunoblot analysis as indicated (upper panels). # indicates nonspecific bands. C, D Immortalized PrxII KO MEFs that had been transfected with expression vectors for PrxII (C172S) or PrxII (WT) or with the corresponding empty vector were incubated for 30 min in the absence or presence of GOX (0.5 or 5 mU/ml) (C) or H_2_O_2_ (50 or 250 μM) (D), after which cell lysates were subjected to nonreducing SDS-PAGE followed by immunoblot analysis as indicated (upper panels). The relative ratio of oxidized (disulphide) HMGB1 to total HMGB1 was estimated by measurement of band intensities in each immunoblot (lower panels). Data are means ± SEM for five (A, B) or three (C, D) independent experiments. \**P* < 0.05, \*\**P* < 0.01 (Student’s *t* test) versus corresponding value for PrxII KO cells (A, B) or for cells transfected with the empty vector (C, D). See also Expanded View figure 2.

Taken together, our data indicated that (1) both PrxI and PrxII sense H_2_O_2_ and form a disulphide-linked homodimer for transduction of the H_2_O_2_ signal by transfer of their oxidation state to HMGB1, thereby generating disulphide HMGB1; (2) the transfer of oxidation state from the Prx dimer to reduced HMGB1 involves formation of a Prx-HMGB1 complex linked by an intermolecular disulphide between the C_P_ residue of Prx and either Cys^23^ or Cys^45^ of HMGB1; and (3) the intermolecular disulphide is resolved by the cysteine residue (Cys^23^ or Cys^45^) of HMGB1 not contributing to the intermolecular linkage, resulting in the production of disulphide HMGB1. It is of note that HMGB1 oxidation occurs in both PrxI and PrxII KO MEFs, albeit a reduced rate compared with that in WT cells. This is likely because of the redundant function of PrxI and PrxII. It was not possible to knock out both Prx isoforms because of the lethality of thus manipulation in mice.

### Oxidation to the disulphide form is necessary for nucleocytoplasmic translocation and secretion of HMGB1

We examined the relevance of the oxidation of HMGB1 to the disulphide form with regard to its nucleocytoplasmic translocation and secretion in HEK293T cells stably transfected with 10 an expression plasmid for TLR4 (HEK293T-TLR4 cells). These cells were further transfected with enhanced green fluorescent protein (EGFP)–tagged WT, C23S, or C45S forms of HMGB1. LPS induced nucleocytoplasmic translocation of EGFP-tagged HMGB1(WT), as revealed by a prominent increase in the level of EGFP fluorescence in the cytoplasm, whereas it had no such effect on EGFP-tagged HMGB1(C23S) or HMGB1(C45S), each of which forms a disulphide-linked complex with PrxI/II that cannot be further resolved to yield disulphide HMGB1 (Fig 3A and B). Immunoblot analysis of culture supernatants revealed that LPS induced the secretion of endogenouse-HMGB1 in MEFs (Fig 3C). The secreted protein was judged to be the disulphide form on the basis of the observation that its mobility on nonreducing SDS-PAGE was decreased by the addition of DTT to the culture supernatants (Fig 3C).

**Figure 3.**
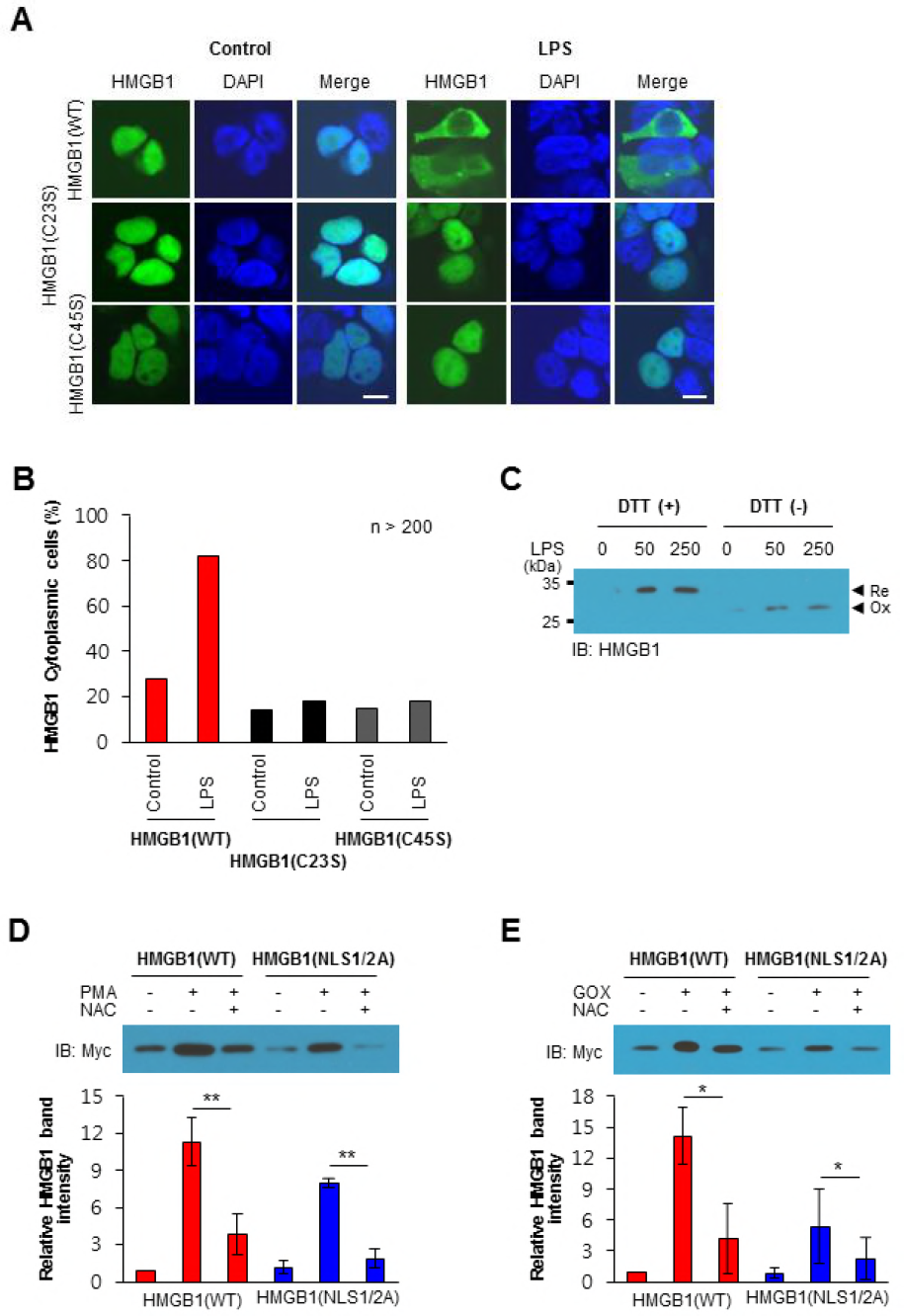
Oxidation of HMGB1 is required for nucleocytoplasmic translocation and secretion. A, B HEK293T-TLR4 cells expressing Myc/EGFP-tagged WT, C23S, or C45S forms of HMGB1 were incubated in the absence (Control) or presence of LPS (1 üg/ml) for 2 h, after which the cells were stained with DAPI and examined for EGFP fluorescence by confocal microscopy (A). Scale bars, 10 üm. EGFP-expressing cells (>200) were scored for the presence of cytoplasmic EGFP, and the percentage of positive cells was determined (B). C. MEFs were stimulated with LPS (50 or 250 ng/ml) for 24 h, after which culture supernatants were harvested, mixed or not with DTT, and subjected to nonreducing SDS-PAGE followed by immunoblot analysis with antibodies to HMGB1. D, E HEK293T cells expressing Myc-HMGB1(WT) or Myc-HMGB1(NLS1/2A) were incubated in the absence or presence of 10 mM NAC or 250 nM PMA for 24 h (D) or of 10 mM NAC or GOX (50 μU/ml) for 24 h (E). Culture supernatants were then harvested and subjected to immunoblot analysis with antibodies to Myc (upper panels). Relative Myc-HMGB1 band intensities were determined as means ± SEM from three independent experiments (lower panels). \**P* < 0.05, \*\**P* < 0.01 (Student’s *t* test). See also Expanded View figure 3.

We also examined the nucleocytoplasmic transport toward secretion of Myc/EGFPHMGB1 induced by H_2_O_2_ or GOX in HEK293T cells. Treatment with H_2_O_2_ or GOX induced nucleocytoplasmic translocation (Fig EV3A and B) as well as extracellular secretion (Fig EV3C and D) of Myc-HMGB1(WT) but not of Myc-HMGB1(C23S) or Myc-HMGB1(C45S),suggesting that disulphide bond formation between Cys^23^ and Cys^45^ is also required for H_2_O_2-_induced HMGB1 translocation and secretion.

Like HMGB1 phosphorylation in PMA-treated cells, acetylation of the NLS sequences of HMGB1 in cells treated with the histone deacetylase inhibitor TSA promotes the nuclear transport of HMGB1 toward secretion (Bonaldi et al, 2003; Youn & Shin, 2006). Whereas PKC activation by PMA results in H_2_O_2_ production through Nox activation (Karlsson et al, 2000), TSA-induced acetylation of multiple mitochondrial proteins increases mitochondrial H_2_O_2_ production (Sun et al, 2014). Binding of TNFα to its cognate receptor induces both acetylation and phosphorylation of the NLS sequences of HMGB1 and thereby promotes HMGB1 secretion (Bonaldi et al, 2003; Willenbrock et al, 2012; Youn & Shin, 2006). TNFα binding also induces H_2_O_2_ production from multiple sources including mitochondria, Nox, and xanthine oxidase (Chen et al, 2008). Treatment of MEFs or HEK293T cells with PMA, TSA, or TNFα stimulated HMGB1 release in a manner sensitive to inhibition by the antioxidant *N*-acetylcysteine (NAC) (Fig EV3E and F), suggesting that oxidant production plays a key role in HMGB1 secretion. The Nox inhibitor diphenyleneiodonium (DPI) inhibited HMGB1 secretion induced by PMA but not that induced by TNFα (Fig EV3G), suggesting that H_2_O_2_ produced by Nox is required for PMA-induced HMGB1 secretion whereas non-Nox sources of H_2_O_2_ such as mitochondria and xanthine oxidase are important for HMGB1 secretion induced by TNFα. Given that H_2_O_2_affects many signaling pathways by targeting proteinaceous cysteine thiols, we examined whether the oxidation of Cys^23^ and Cys^45^ is also required for the nuclear transport toward secretion of HMGB1 in response to PMA, TSA, or TNFα. All three stimulants triggered the nucleocytoplasmic translocation (Fig EV3H and I) and secretion (Fig EV3J-L) of Myc-HMGB1(WT) but not those of Myc-HMGB1(C23S) or Myc-HMGB1(C45S). These results suggested that disulphide bond formation between Cys^23^ and Cys^45^ of HMGB1 is a prerequisite for its secretion whereas acetylation or phosphorylation is not sufficient.

We next tested whether a phosphorylation-defective mutant (NLS1/2A, in which all six serines in the NLS sequences are replaced with alanine) of HMGB1 (Youn & Shin, 2006) is secreted in response to oxidative stimuli. Stimulation of HEK293T cells expressing Myc-HMGB1(WT) or Myc-HMGB1(NLS1/2A) with PMA or GOX revealed that secretion of the mutant protein did occur in response to each stimulant but at a level lower than that apparent for the WT protein, with secretion of both HMGB1(WT) and HMGB1(NLS1/2A) being greatly attenuated by NAC (Fig 3D and E).

Collectively, these results indicated that intramolecular disulphide bond formation between Cys^23^ and Cys^45^ of HMGB1 is required for nucleocytoplasmic transport toward secretion in response to all the stimulants tested (LPS, H_2_O_2_, PMA, TSA, and TNFα).Posttranslational modification such as phosphorylation and acetylation of the NLS sequences appears not to be required for HMGB1 translocation toward secretion but rather facilitates this process. PMA thus induced the secretion of an HMGB1 mutant without NLS phosphorylation sites, but the extent of this effect was less pronounced than that apparent with the WT protein. The observation that GOX-induced HMGB1 secretion was also partially inhibited by NLS mutation is likely related to the fact that PKC isoforms are activated by H_2_O_2_ independently of diacylglycerol, the PKC activator mimicked by PMA (Konishi et al,1997) .

### Preferential binding of disulphide HMGB1 to CRM1

HMGB1 is sufficiently small to undergo passive diffusion through nuclear pores. Its nucleocytoplasmic translocation is also actively regulated by the nuclear exportin CRM1, to which HMGB1 binds directly (Bonaldi et al, 2003). We examined the affinity of HMGB1 for CRM1 with the use of HEK293T cells transfected with expression vectors for CRM1 and Myc epitope–tagged WT, C23S, or C45S forms of HMGB1. Immunoprecipitation revealed that CRM1 binding to HMGB1(WT) was greatly increased by exposure of the cells to H_2_O_2_ or GOX (Fig 4A). In the treated cells, the apparent binding affinity of HMGB1 for CRM1 decreased in the rank order of WT > C23S > C45S (Fig 4A). Similar results were obtained by PLA analysis of the proximity of the Myc epitope–tagged forms of HMGB1 and endogenous CRM1 (Fig 4B). Together, these results thus indicated that the affinity of disulphide HMGB1 for CRM1 is greater than that of reduced HMGB1, likely explaining the increased rate of nucleocytoplasmic translocation of the disulphide form observed in the presence of various stimulants.

**Figure 4.**
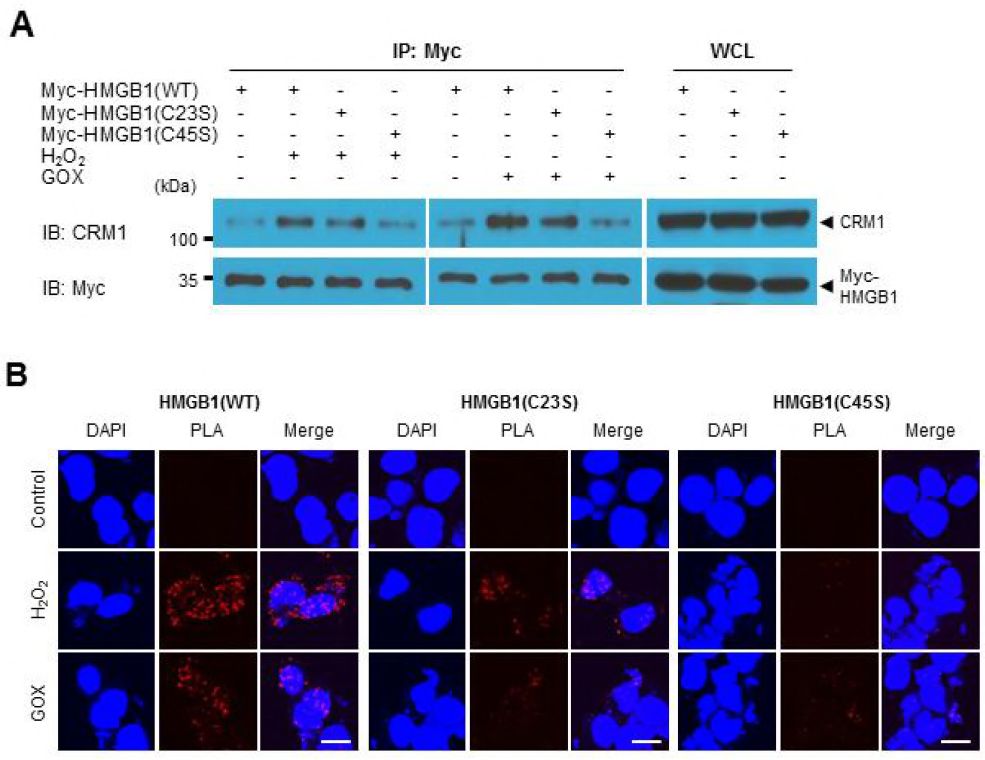
Binding of disulphide HMGB1 to CRM1. A. HEK239T cells transfected with expression vectors for CRM1 and Myc epitope–tagged WT, C23S, or C45S forms of HMGB1 were incubated in the absence or presence of 50 üM H_2_O_2_ or GOX (0.5 mU/ml) for 2 h, after which cell lysates were subjected to immunoprecipitation with antibodies to Myc. The resulting precipitates as well as the whole cell lysates (WCL) were subjected to immunoblot analysis with antibodies to CRM1 and to Myc. B. HEK293T cells expressing Myc epitope–tagged WT, C23S, or C45S forms of HMGB1 were incubated in the absence or presence of 50 μM H_2_O_2_ or GOX (5 mU/ml) for 2 h and then subjected to a PLA assay with antibodies to Myc and to CRM1. Scale bars, 10 üm.

### HMGB1 secretion from BMDMs is dependent on PrxI and PrxII

We have shown that HMGB1 oxidation to the disulphide form is mediated by PrxI and PrxII and that intramolecular disulphide formation is necessary for nucleocytoplasmic translocation and secretion of HMGB1 in nonimmune cell types such as HEK293T cells and MEFs. We therefore next investigated the role of PrxI and PrxII in HMGB1 secretion with the use of PrxI- or PrxII-deficient BMDMs (Fig EV4A). HMGB1 secretion induced by LPS, PMA, TSA, TNFα, H_2_O_2_, or GOX was attenuated in PrxI- or PrxII-deficient BMDMs compared with WT BMDMs (Fig 5; Fig EV4B and C). As was the case for intramolecular disulphide formation in nonimmune cells (Fig 2A and B), the suppression of HMGB1 secretion by PrxI or PrxII deficiency in BMDMs was partial. Partial inhibition of HMGB1 secretion induced by LPS or GOX was also apparent in PrxII KO MEFs compared with WT MEFs, whereas the extent of HMGB1 secretion was fully restored in the mutant cells by forced expression of PrxII (Fig EV4D and E).

**Figure 5.**
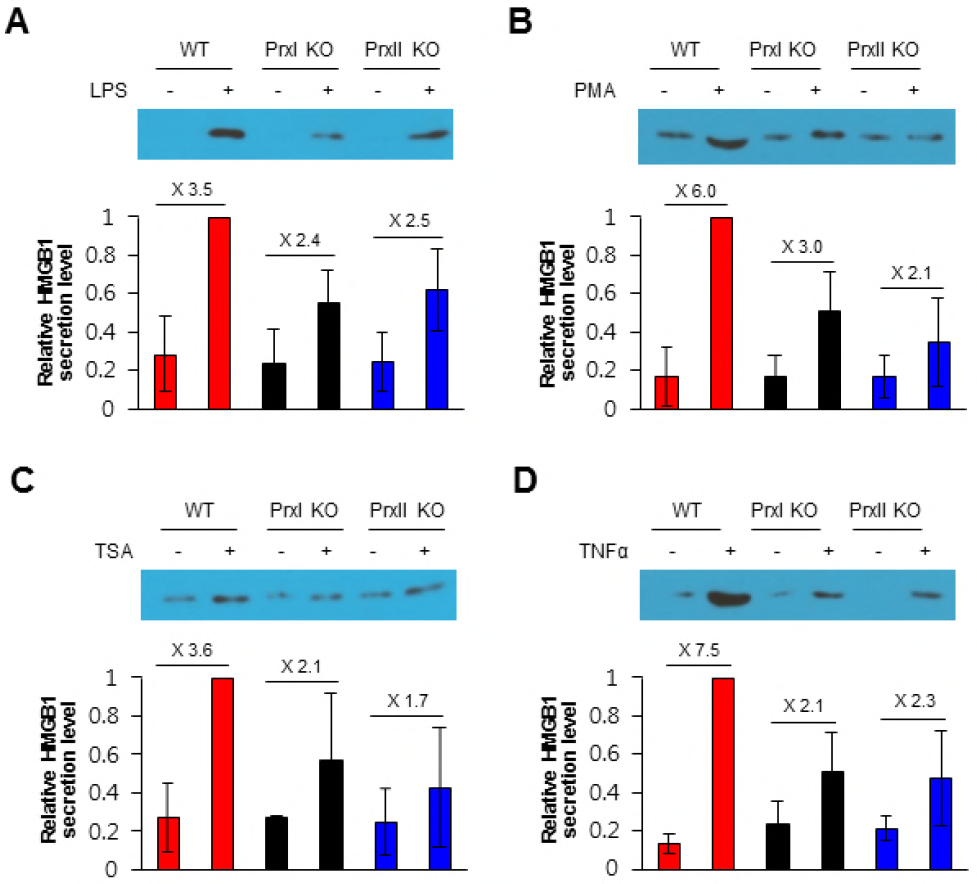
Deficiency of PrxI or PrxII attenuates HMGB1 secretion in BMDMs. BMDMs derived from WT, PrxI-deficient (PrxI^−/−^), or PrxII-deficient (PrxII^−/−^) mice were incubated in the absence or presence of LPS (100 ng/ml) (A), 250 nM PMA (B), TSA (50 ng/ml) (C), or TNFα (50 ng/ml) (D) for 24 h, after which culture supernatants were subjected to immunoblot analysis with antibodies to HMGB1 (upper panels). Relative HMGB1 band intensities (secretion levels) are presented as means ± SEM from three independent experiments (lower panels). The fold induction of secretion by each stimulus is also shown. See also Expanded View figure 4.

### LPS-induced HMGB1 secretion is dependent on PrxI and PrxII in vivo

Finally, we examined the role of PrxI and PrxII in HMGB1 secretion in vivo. WT as well as PrxI- or PrxII-deficient mice were injected intraperitoneally with a sublethal dose (5 mg/kg) of LPS, and blood samples were collected at optimal times (1 and 16 h, respectively) for the measurement of TNFα and HMGB1 concentrations in serum. LPS injection increased TNFα levels to 22.0 ± 6.7 ng/ml in WT mice, to 15.7 ± 6.1 ng/ml in PrxI-deficient mice, and to 24.3 ± 6.9 ng/ml in PrxII-deficient mice (Fig 6A), values that did not differ significantly from each other. In contrast, LPS increased HMGB1 levels in PrxI- or PrxII-deficient mice to 31.4 ± 13.6 and 43.6 ± 22.6 ng/ml, respectively, values that were significantly smaller than the corresponding value of 74.2 ± 14.7 ng/ml for WT mice (Fig 6B).

**Figure 6.**
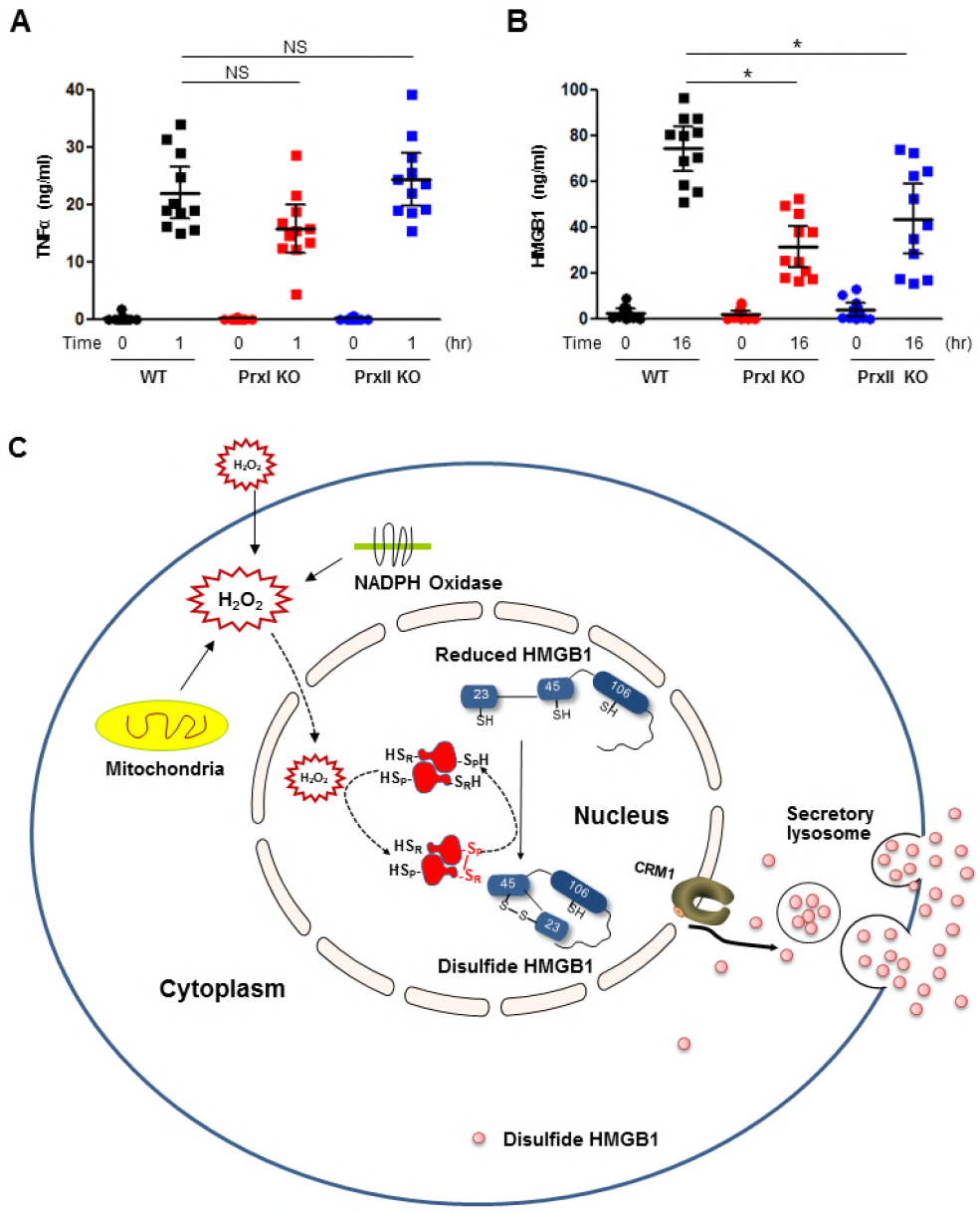
Serum HMGB1 levels in PrxI- or PrxII-deficient mice injected with LPS. A, B WT, PrxI^−/−^, or PrxII^−/−^ mice (*n* = 11 per group) were injected intraperitoneally with LPS (5 mg/kg), and blood samples were collected for preparation of serum at 1 and 16 h, corresponding to the peak levels of TNFα and HMGB1, respectively. The concentrations of TNFα (A) and HMGB1 (B) in serum were measured with enzyme-linked immunosorbent assay kits. Data for individual mice as well as mean ± SD values for each group are shown. \**P* < 0.01 (Student’s *t* test). C. Model for the mechanism underlying HMGB1 secretion mediated by PrxI and PrxII in response to various inflammatory stimuli.

## Discussion

The nuclear protein HMGB1 passively leaks from necrotic and apoptotic cells of various types and is actively secreted from live monocytes-macrophages in response to stress signals (Kang et al, 2014). HMGB1 in the extracellular milieu serves as a DAMP molecule. Posttranslational modification such as acetylation of lysine residues or phosphorylation of serine residues located in the NLS sequences of HMGB1 has been proposed as the major mechanism for regulation of active nuclear exit of the protein (Bonaldi et al, 2003; Youn & Shin, 2006). All stimulants of HMGB1 secretion tested in the present study (LPS, TNFα, PMA, and TSA) are known to induce the intracellular production of H_2_O_2_ at low levels, with oxidation of various proteinaceous cysteine residues by H_2_O_2_ being an important mechanism of cell signalling (Poole & Nelson, 2008). Necrotic and apoptotic cell death is also accompanied by H_2_O_2_ production by mitochondria (Circu & Aw, 2010; Fiers et al, 1999).

In the present study, we found that oxidation of Cys^23^ and Cys^45^ to yield disulphide HMGB1 is required for nucleocytoplasmic translocation and secretion of the protein. The C23S and C45S mutants of HMGB1, which are not able to form the intramolecular disulphide bond but in which acetylation and phosphorylation sites are intact, thus manifested almost undetectable levels of nucleocytoplasmic translocation and secretion in response to various stimulants. The dispensability of NLS phosphorylation for HMGB1 secretion was also demonstrated by the observation that the phosphorylation-defective NLS1/2A mutant was still secreted in response to H_2_O_2_ stimulation. Despite the facts that oxidation of protein thiols is generally a slow process and that cellular compartments are equipped with various H_2_O_2_-eliminating enzymes, the specific oxidation of HMGB1 by H_2_O_2_ at low concentrations can be achieved in the nucleus as a result of the H_2_O_2_ sensor and transducer function of PrxI and PrxII. Such a function of PrxI and PrxII in the case of HMGB1 was demonstrated by direct detection of the reaction intermediates comprising complexes of HMGB1 and either PrxI or PrxII linked by an intermolecular disulphide between the C_P_ residue of PrxI/II and either Cys^23^ or Cys^45^ of HMGB1.

On the basis of our findings, we propose a model for the mechanism underlying regulated HMGB1 secretion from monocyctes and macrophages. Inflammatory stimuli such as LPS and TNFα as well as chemical agents such as PMA and TSA induce H_2_O_2_ production from multiple sources including Nox and mitochondria in the cytoplasm, and the H_2_O_2_ molecules than diffuse to the nucleus. It is also likely that H_2_O_2_ produced as a result of the respiratory burst of neighboring neutrophils and macrophages can enter the cell and reach the nucleus, and that activation of Nox enzymes localized at the nuclear membrane may be a source of nuclear H_2_O_2_ (Provost et al, 2010; Stanicka et al, 2015). The accumulation of H_2_O_2_ in the nucleus is first sensed by PrxI and PrxII, resulting in the rapid oxidation of C_P_ and C_R_ of each enzyme and the consequent formation of intersubunit disulphide bonds in each dimer.The oxidized PrxI or PrxII then transfers its oxidation state to reduced HMGB1 to generate disulphide HMGB1 with an intramolecular disulphide between Cys^23^ and Cys^45^. Disulphide HMGB1 is preferentially transported out of the nucleus as a result of its binding to the nuclear exportin CRM1 with higher affinity compared with that of reduced HMGB1— although, given that disulphide HMGB1 is more compact than the reduced form, preferential passive diffusion of disulphide HMGB1 cannot be excluded. It is also possible that, like acetylated HMGB1, disulphide HMGB1 may be less favoured for nuclear import and thus accumulates in the cytosol. Disulphide HMGB1 in the cytosol is packed into lysosomes through an as yet unknown mechanism and is then secreted. Our results show that HMGB1 that is actively secreted in response to LPS stimulation is present in the disulphide form, not the reduced form. Given that the cytosol contains various disulphide reductases such as thioredoxins and glutaredoxins that are capable of mediating the rapid reduction of disulphide HMGB1 to reduced HMGB1 (Hoppe et al, 2006), the H_2_O_2_ sensor and transducer function of PrxI and PrxII might also be needed in the cytosol to maintain HMGB1 in the disulphide form before it is packed into lysosomes.

## Materials and Methods

### Antibodies and chemicals

All antibodies for immunoblotting analysis were purchased from Abcam, Santa Cruz Biotechnology, Invitrogen Life Technologies, and AbFrontier Chemicals including LPS (L5293, Sigma), trichostatin A (T8552, Sigma), phorbol 12-myristate 13-acetate (P8139, Sigma), and *N*-ethyl maleimide (E3876, Sigma) were purchased from commercial venders.

### Cell cultures

HEK293T, HEK293T-TLR4, and MEF cells were cultured in 10% FBS-Dulbecco’s modified Eagle’s medium (DMEM) supplemented with 100 U/mL penicillin, 100 üg/mL streptomycin, and 2 mM L-glutamine at 37°C under 5% CO_2_. Primary mouse BMDM cells were prepared from male C57BL/6 WT, PrxI-, and PrxII-deficient mice (Bae et al, 2011; Lee et al, 2003). Briefly, bone marrow cells were obtained by flushing bone marrow of femurs and tibias after euthanasia and then maintained in complete DMEM medium containing 20% L929 culture supernatant for 7 days (Weischenfeldt & Porse, 2008). BMDM cells were collected after incubation at 37°C for 4 min with 0.25% trypsin-EDTA solution and were gently resuspended in complete DMEM.

### Animals

C57BL/6 WT, PrxI-, and PrxII-deficient mice were used (Bae et al, 2011; Lee et al, 2003). Mice were performed on age- and gender-matched randomly assigned 8- to 12-week-old male mice conducted according to procedures approved by the Institutional Animal Care and Use Committee of the Yonsei Laboratory Animal Research Center (YLARC, 2015-0275). Up to five mice per cage were housed in ventilated cages in a 12 hr light and 12 hr dark cycle. Room temperature was at 22°C, and they were fed with autoclaved standard rodent food and with drinking water *at libitum*.

### Plasmids and in situ mutagenesis

Myc- or enhanced green fluorescent protein (EGFP)-tagged HMGB1^WT^, HMGB1^C23S^, and HMGB1^C45S^ were inserted into the pCMV-Myc plasmid for mammalian cell expression. Myc/His-tagged human PrxI and PrxII plasmids were generated using the pCDNA3.1 vector. Myc/His-tagged PrxI^C52S^, PrxI^C173S^, PrxI^C52S/C173S^, PrxII^C51S^, PrxII^C172S^, and PrxII^C51S/C172S^ plasmids were generated using the QuickChange site-directed mutagenesis kit. Streptavidin-binding peptide (SBP)-tagged PrxII^WT^, PrxII^C51S^, PrxII^C172S^, and PrxII^C51S/C172S^ plasmids were kindly provided by Dr. Tobias P. Dick (German Cancer Research Center, Heidelberg, Germany) (Sobotta et al, 2015), and subcloned into pcDNA^TM^3.1/Myc-His. The HMGB1^NLS1/2A^-GFP plasmid, a mutant plasmid wherein all the phosphorylation residues of six serines are changed to alanines in the NLS1 and 2 regions (Youn & Shin, 2006), was subcloned into pCMV-Myc.

### Cell culture, transfection, and reagents

Plasmid transfections were carried out using Fugene HD reagent or electroporation by MicroPorator-mini. The histone deacetylase inhibitor TSA and PKC activator PMA for acetylation and phosphorylation, respectively, as well as H_2_O_2_ and GOX from *Aspergillus niger* for oxidation were used. LPS (*Escherichia coli* 0111:B4, 49180) and TNF-α were used for pro-inflammatory signaling. NAC was used as an antioxidant.

### Measurement of HMGB1 secretion

To analyze the secretion of HMGB1 in the supernatants, culture media were replaced with serum-free OPTI-MEM medium and treated with various stimulants. The culture supernatants were harvested and concentrated via Amicon Centricon filtration after removing cell debris. Western blotting analysis was performed with rabbit anti-HMGB1 or mouse anti-Myc antibodies as described below.

### In vitro binding of PrxI/II proteins to HMGB1

HMGB1 and PrxI or PrxII proteins were reduced with 5 mM DTT for 1 hr and then the DTT was removed before the in vitro binding assay. A mixture of HMGB1 and PrxI or PrxII was exposed to 50 üM H_2_O_2_ with 0, 5, or 50 üM DTT for 30 min to observe disulphide bond formation. The mixture was immunoprecipitated with mouse monoclonal anti-HMGB1 Ab and immunoblotted with rabbit polyclonal anti-PrxI, rabbit monoclonal anti-PrxII or rabbit polyclonal anti-HMGB1 antibodies after washing.

### Confocal microscopy

Nucleocytoplasmic translocation of HMGB1 was observed under various stimulation conditions. HEK293T and HEK293T-TLR4 cells were transfected with EGFP-tagged HMGB1^WT^, HMGB1^C23S^, and HMGB1^C45S^ plasmids and cultured in a LabTekTM II chamber for 48 hr. Cells were treated with 5 mU/mL GOX, 50 μM H_2_O_2_, 250 nM PMA, 50 ng/mL TNF-α, and 10 ng/mL TSA for 2 hr. HEK293T-TLR4 cells were treated with 1 üg/mL L for 1 hr. Cells were fixed with 4% paraformaldehyde in PHEM buffer for 20 min at room temperature (RT) and washed with cold PBS. After mounting with 4’,6’-diamidino-2-phenylindole, over 200 cells were observed to count the proportion of cytoplasmic HMGB1 (+) cells under confocal FV1000 microscopy (Olympus).

### Proximity ligation assay (PLA)

The molecular interactions between HMGB1 and PrxI/II or CRM1 proteins were evaluated using a Duolink II Detection kit (Olink Bioscience). Briefly, HEK293T cells were cultured in eight well chambers (Nunc) and transfected with HMGB1^WT^, HMGB1^C23S^, or HMGB1^C45S^ plasmids. Cells were fixed with 4% paraformaldehyde in PHEM buffer for 20 min at RT. Cells were incubated with a blocking agent for 1 hr, and rabbit polyclonal anti-PrxI and anti-PrxII were added and incubated overnight with either mouse monoclonal anti-HMGB1 or anti-myc for endogenous or exogenous HMGB1, respectively. Mouse monoclonal anti-CRM1 was incubated with rabbit polyclonal anti-HMGB1 to identify the HMGB1-CRM1 interaction. After three subsequent washes, PLA probes were applied and incubated for 1 hr in a humidity chamber at 37°C. Unbound PLA probes were removed and the samples were incubated in the ligation solution for 1 hr. For amplification, polymerase was applied with fluorescently labeled oligonucleotides for 100 min at 37°C. The amplified fluorescence-labeled oligonucleotide was visible as a distinct fluorescent spot and images were taken by confocal microscopy.

### Mouse study

C57BL/6 WT, PrxI- and PrxII-deficient mice were used to investigate the effect of PrxI/II on HMGB1 secretion in vivo. Mice were intraperitoneally injected with 5 mg/kg LPS and then serum samples were collected at 1 and 16 hr for TNF-α and HMGB1 measurements, respectively. Serum HMGB1 and TNF-α were measured according to the manufacturer’s protocols.

### Western blot analysis and immunoprecipitation

HEK293T or MEF cells in six-well plates were transfected with plasmids using FuGene HD or electroporated using the MicroPorator-mini. Cells were washed with phosphate buffered saline (PBS) and lysed in 1× RIPA buffer containing 150 mM NaCl, 1% Triton X-100, 1% deoxycholic acid sodium salt, 0.1% SDS, 50 mM Tris-HCl pH 7.5, 2 mM EDTA, and protease inhibitor cocktail. For thiol blocking, 50 mM NEM was added to inhibit the breakage or new disulphide bond formation. Whole cell lysates (WCLs) were centrifuged at 20,000 × *g* for 10 min at 4°C. Protein sample buffer (100 mM Tris-HCl pH 6.8, 2% SDS, 25% glycerol, 0.1% bromophenol blue, and 5% β-mercaptoethanol) was added to the WCLs followed by heating at 94°C for 5 min. Proteins (20 μg) were separated by molecular weight via SDS-PAGE and transferred to nitrocellulose membranes . Non-specific binding sites were blocked by incubating the membranes in Tris-buffered saline (TBS) supplemented with 0.1% Tween 20 and 5% w/v skim milk for 1 hr. Anti-HMGB1, anti-PrxI, anti-PrxII, anti-CRM1, anti-GAPDH, anti-Sp1, and anti-Myc antibodies were used. Membranes were washed three times for 10 min with TBS-Tween 20 and probed with the appropriate horseradish peroxidase (HRP)-conjugated secondary antibody for 1 hr. After washing three times, an enhanced chemiluminescence substrate was then used for visualization. Membranes were then stripped by submerging in EzReprobe at room temperature for 30 min.

Immunoprecipitation was performed with 100–200 μg protein. Initially, 1 üg specific antibody was conjugated with Dynabeads^^®^^ protein G at room temperature for 1 hr. Samples were added to 10 μL antibody conjugated-Dynabeads^^®^^ Protein G at 4°C for 18 hr. The beads were washed three times with ice-cold PBS and then mixed with protein sample buffer followed by heating at 94°C for 5 min. The proteins were separated using SDS-PAGE.

### Measurement of PrxI/II concentration

To determine the concentration of PrxI and PrxII proteins in the nucleus, nuclear fractionation was obtained using nuclear/cytosolic fractionation kit according to the manufactures’ procedure. Fifteen micrograms of each nuclear fraction were separated with known amounts of standard PrxI and II proteins from mouse liver lysate. The concentration of standard PrxI or II protein was previously identified by comparing the band intensities with those of recombinant PrxI and PrxII protein using SDS-PAGE. Anti-PrxI and anti-PrxII antibodies were used for western blot. PrxI and PrxII concentrations were calculated by comparing their band intensities with those of standard protein using Image J.

### Nuclear/cytosolic fractionation

To determine the localization of the intracellular interaction between HMGB1 and PrxI or PrxII during conditions of increased intracellular ROS levels, HEK293T cells were transfected with Myc tagged HMGB1^WT^, HMGB1^C23S^, or HMGB1^C45S^ with PrxI- or PrxIIMyc/His plasmids and then treated with 50 üM H_2_O_2_ for 30 min to induce oxidative stress. Cells were harvested by centrifugation at 600 × *g* for 5 min at 4°C. Nuclear/cytosolic fractionation was performed using a nuclear/cytosolic fractionation kit according to the manufacturer’s procedure.

### Statistical analysis

Analysis of experimental data was performed with the Student’s t test using GraphPad Prism. Data represent the mean value and SEM or SD, as indicated in the individual figure legends. The difference was considered statistically significant when p < 0.05.

## Acknowledgments

We thank Dr. Tobias P. Dick (German Cancer Research Center, Heidelberg, Germany) for providing Prx mutant plasmids (wild type PrxII-SBP and mutants PrxII-SBP C51S, C172S, C52/172S), and Dr. In Sup Kil for helpful discussion. This work was supported by the National Research Foundation of Korea (NRF) funded by the Korean government (MEST) [2014R1A4A1008625 and 2017R1A2B3006704], and by the Research Center Program of Institute for Basic Science (IBS) in Korea [IBS-R026-D1].

## Author contributions

M.S.K. participated in and coordinated the majority of the experiments. H.S.K. and M.G.H. performed PrxI mutant plasmid generation and functional analysis. K.L. performed part of the confocal microscopy analysis. Y.H.K. carried out the interaction between HMGB1 and PrxI/II in the nucleus. J.M.S. analyzed the oxidation levels of HMGB1, PrxI, and PrxII in the nuclei of BMDM cells. I.H.P. participated in the mouse serum study. S.K.L. helped with the mouse study and PrxI/II concentration measurement. S.G.R. participated in study design and discussion, and wrote the manuscript. J.S.S. supervised and coordinated all the experiments and wrote the manuscript. All authors read and approved the final manuscript.

